# An extensive evaluation of single-cell RNA-Seq contrastive learning generative networks for intrinsic cell-types distribution estimation

**DOI:** 10.1101/2025.09.15.675691

**Authors:** Ibrahim Alsaggaf, Daniel Buchan, Cen Wan

**Affiliations:** School of Computing and Mathematical Sciences, Birkbeck, University of London, United Kingdom; Department of Computer Science, University College London, United Kingdom

## Abstract

Contrastive learning has already been widely used to handle single-cell RNA-Seq data due to its outstanding performance in transforming original data distributions into hypersphere feature spaces. In this work, we conduct a large-scale empirical evaluation to investigate the generative encoder networks that are learned by five different state-of-the-art single-cell RNA-Seq contrastive learning methods. Unlike the conventional discriminative model-based cell-type prediction studies, this work is focused on the performance of contrastive learning-based generative encoder networks in terms of their capacity to estimate the intrinsic distributions of different cell-types – a fundamental property that directly affects the performance of any downstream single-cell RNA-Seq data analytics. The experimental results confirm that supervised contrastive learning-based encoder networks lead to better performance than self-supervised contrastive learning-based encoder networks, and the recently proposed Gaussian noise augmentation-based single-cell RNA-Seq contrastive learning method shows the best performance on estimating the intrinsic distribution of different cell-types.

## Introduction

Single-cell RNA sequencing (scRNA-Seq) is an efficient approach for revealing complex transcriptome landscapes at single-cell resolution. Due to the abundant transcriptome information, scRNA-Seq plays a crucial role in understanding sophisticated biological processes and has achieved great success in a variety of important biological challenges, such as cell-type identification [1–3], gene regulatory mechanism prediction [4–6], and drug discovery [7–9]. However, one of the fundamental challenges of scRNA-Seq analytics is dealing with the high-dimensional distributions of transcriptome landscapes. Due to the large number of genes included in cells, the dimensionality of scRNA-Seq data is usually extremely high. For example, the well-known peripheral blood mononuclear cells (PBMCBench 10Xv2) [3, 10] dataset includes 22,280 genes’ expression profiles for 6,444 cells. Such high-dimensional data inevitably leads to natural difficulties in accurately estimating the high-dimensional distributions of scRNA-Seq data using computational methods due to the well-known curse of dimensionality issues.

Contrastive learning is an emerging machine learning paradigm that aims to learn a hypersphere feature representation space where similar instances are pulled closer and dissimilar instances are pushed away from each other. The embeddings derived from the learned hypersphere space encode discriminative information for downstream predictive tasks. For example, in computer vision and natural language processing research, contrastive learning-based predictive models marginally improved the performance of object detection [11–13] and image-text generation [14–16] tasks. Contrastive learning has also been widely used in bioinformatics research recently, such as protein–DNA binding sites prediction [17], protein function prediction [18] and ageing-related gene effects prediction [19]. In terms of scRNA-Seq analytics tasks, contrastive learning has also demonstrated superb performance. For example, Ciortan and Defrance (2021) [20] first proposed a random gene masking augmentation-based contrastive learning method for cell clustering tasks. Yan, et al. (2022) [21] proposed a contrastive learning-based framework for scRNA-Seq integration and batch effects removal tasks. More recently, Alsaggaf, et al. (2024, 2025) [1, 22] proposed a Gaussian noise augmentation-based and an augmentation-free contrastive learning methods that obtained state-of-the-art performance in cell-type identification tasks. However, the capacity of contrastive learning methods for estimating intrinsic cell-type distributions is still understudied. Unlike the conventional discriminative model-based cell-type identification studies [3] that aim to learn decision boundaries of different cell-types, in this paper, we investigate five types of state-of-the-art scRNA-Seq contrastive learning methods in terms of their generative encoder networks’ performance on learning the intrinsic distributions of different cell-types.

This paper is organised as follows. The methods section introduces the proposed contrastive learning generative encoder networks evaluation approach. The computational experiments section reports experimental results, followed by the discussion section where further analysis of different contrastive learning generative encoder networks is reported. Finally, the conclusion section summarises the findings of this work and presents future research directions.

## Methods

In this work, we propose a new method, namely scRCL-G, to evaluate the performance of scRNA-Seq contrastive learning generative encoder networks. In general, the essence of the proposed scRCL-G method aims to evaluate the capacity of contrastive learning-based generative encoder networks to estimate the intrinsic distributions of different cell-types using scRNA-Seq data. More specifically, we aim to investigate the quality of generative encoder networks from two perspectives as shown in the bullet points below.

- The estimated population distributions of cell-types.
- The estimated population centroids of cell-types.

Therefore, in general, scRCL-G uses a training scRNA-Seq dataset (as a sample) to estimate different cell-types’ population characteristics, which are evaluated by using a testing scRNA-Seq dataset. The target of the proposed scRCL-G method is also different from both the discriminative model-based cell-type identification tasks and the conventional scRNA-Seq clustering tasks. The conventional discriminative model-based cell-type identification tasks aim to learn decision boundaries for separating different cell-types in a target dataset, but scRCL-G aims to directly learn distributions for all different cell-types. The conventional clustering tasks aim to discover cell clusters included in a target dataset without exploiting any prior knowledge about cell-types, but scRCL-G aims to transform a target scRNA-Seq dataset into a hypersphere space by using a trained encoder that has already learned the distributions of different cell-types.

The flowchart of the proposed scRCL-G method is shown in Figure 1, whilst the high-level pseudocode is shown in Algorithm 1. Figure 1.A illustrates the conventional self-supervised scRNA-Seq contrastive learning framework, where an encoder network and a projection head network are both trained by using pairs of views created for each individual training instance. Analogously, in line 1 of Algorithm 1, an scRNA-Seq contrastive learning model ℱ that consists of an encoder network ℰ and a projection head network 𝒫 is first trained using 𝒮_*a*_, which denotes an input training sample. As shown in Figure 1.B and lines 2 - 3 of Algorithm 1, the trained encoder network ℰ is then used to transform the entire 𝒮_*a*_, and the transformed sample 𝒮_*a*_′ is used to learn the centroids 𝒞 of different cell-types by conducting the well-known k-means clustering operation. Note that the value of *k* is pre-defined according to the cell-type label information included in 𝒮_*a*_, i.e. *k* equals the total number of cell-types. Lines 4 - 6 of Algorithm 1 show the testing stage of scRCL-G. In line 4, the testing dataset 𝒮_*b*_, which is another input as shown by the white dots in Figure 1.B, is transformed into 𝒮_*b*_′ by using ℰ. In line 5, the learned centroids 𝒞 are applied to the transformed testing sample 𝒮_*b*_′ to obtain cell groups 𝒢, as shown by the dots in different colours denoting assigned cell-types according to the shortest distances between cells and different centroids.

**Fig 1.**
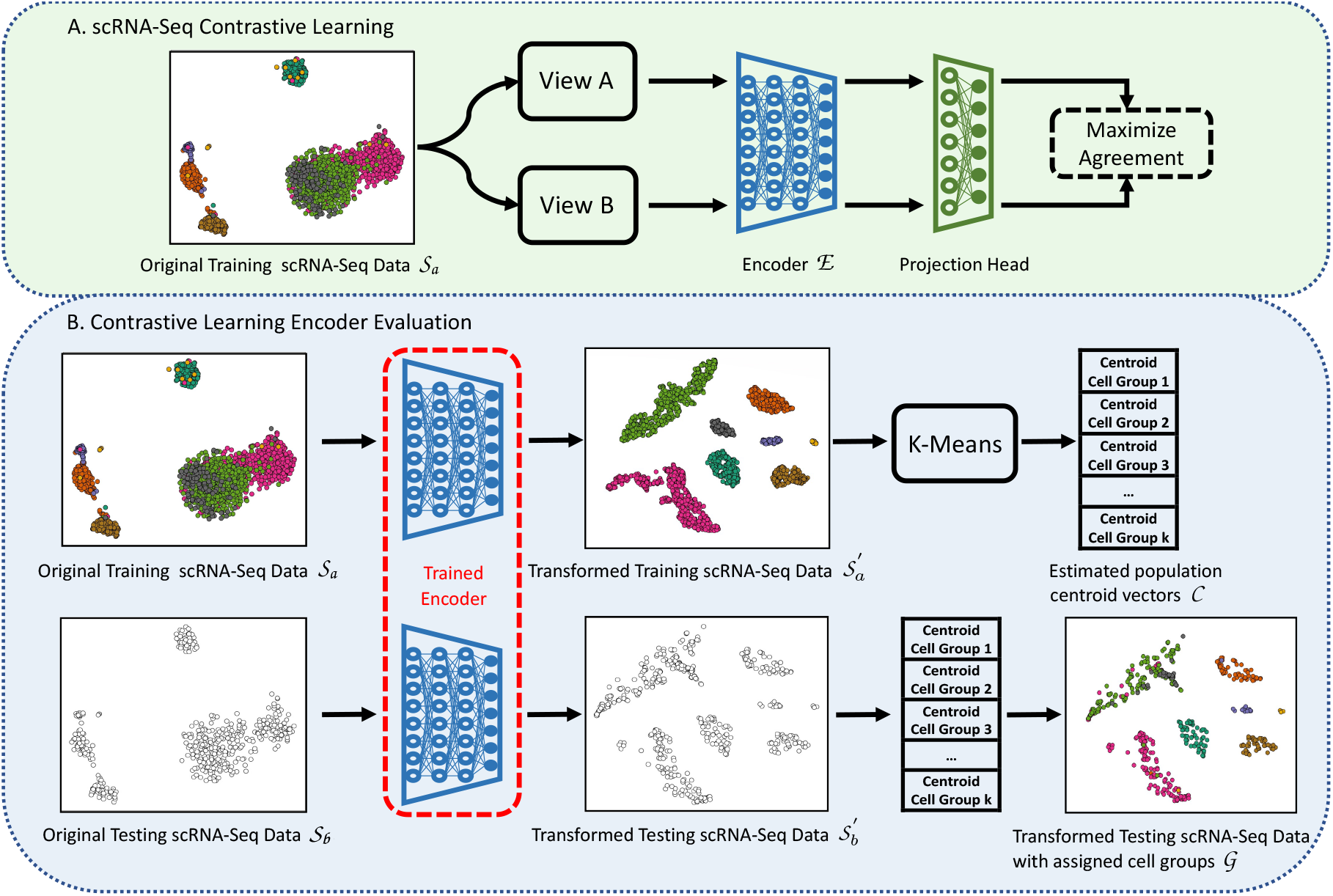
The flowchart of the proposed scRNA-Seq contrastive learning generative encoder evaluation methods (scRCL-G). The different colours (except white) denote different cell types. The dots in white colour denote the cells with unknown cell types.

Finally, in line 6, the quality of ℰ is evaluated by different metrics ℳ with considering the transformed 𝒮_*b*_′ and the learned 𝒢. The values of the metrics ℳ are returned as outputs of Algorithm 1.

### Algorithm 1

scRNA-Seq contrastive learning encoder evaluation method (scRCL-G).

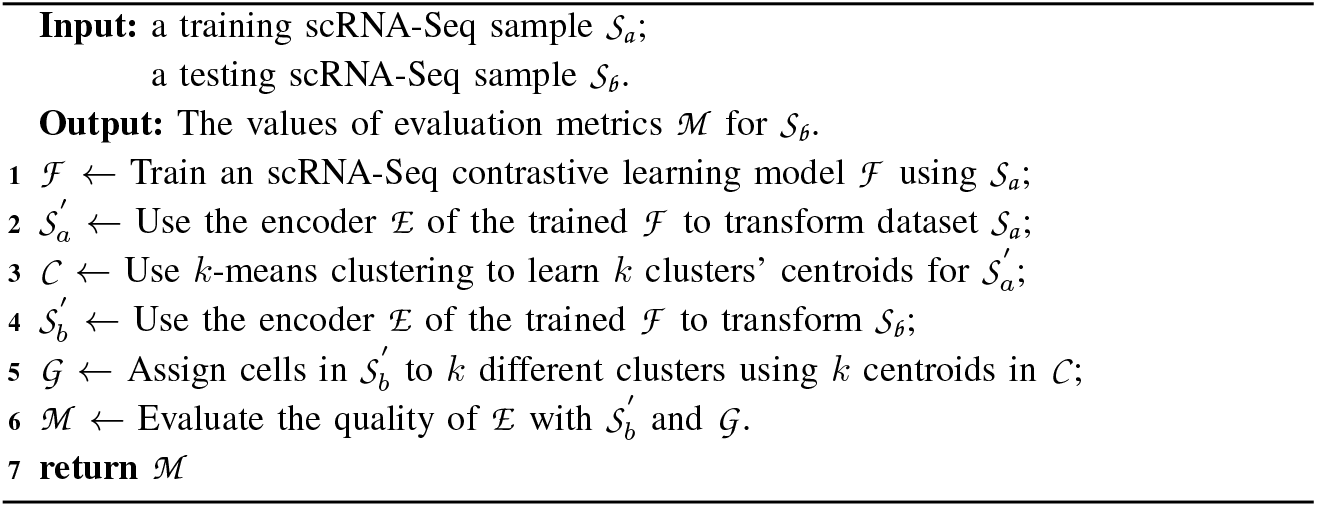

In this work, we use four well-known metrics [23] to evaluate the quality of the predicted cell groups, i.e. Adjusted Rand Index (ARI), Normalized Mutual Information (NMI), Calinski and Harabasz score, and Silhouette score. The higher values of those metrics all mean the better performance of the generative encoder network in terms of its capacity to estimate the distribution of different cell-types. However, note that those metrics do not evaluate the correctness of cell group assignments – they only consider how well different cell groups can be generated.

The ARI score first computes the Rand Index (RI) score, i.e. the agreement between two clusters ignoring permutations, by dividing the number of agreeing pairs over the total number of pairs, then it adjusts the RI score for chance. Let *C*^*t*^ and *C*^*p*^ respectively denote the ground truth labels and the assigned labels, Equation 1 defines the ARI score based on a square contingency matrix, where *TP* is the number of pairs that are in the same cluster in both *C*^*t*^ and *C*^*p*^, *TN* is the number of pairs that are in different clusters in both *C*^*t*^ and *C*^*p*^, *FP* is the number of pairs that are in different clusters in *C*^*t*^ but in the same cluster in *C*^*p*^, and *FN* is the number of pairs that are in the same cluster in *C*^*t*^ but in different clusters in *C*^*p*^.

The NMI score computes the mutual information (MI) between two clusters also ignoring permutations. As shown in the numerator of Equation 2, *k*^*t*^ and *k*^*p*^ respectively denote the number of clusters in the ground truth labels and the assigned labels, whilst 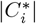 denotes the number of samples in the *i*^*th*^ cluster and *N* denotes the sum of the contingency matrix (i.e. *N* = *TP* + *TN* + *FP* + *FN*). *H* denotes the average of both entropies of *C*^*t*^ and *C*^*p*^ that is used as the denominator to normalise the numerator leading to a value ranging between 0 and 1.

The Calinski and Harabasz score is defined as the ratio of the sum of between-cluster dispersion and within-cluster dispersion without considering the ground truth labels. As shown in Equation 3, |*C*| denotes the total number of instances, and *k* denotes the number of clusters. 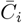 denotes the centroid of the *i*^*th*^ cluster and 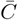denotes the overall centroid.

Analogously, the Silhouette score also does not consider ground truth label information. It computes the Silhouette coefficient for each instance *x* using the average intra-cluster distance and the average distance to the nearest cluster. As shown in Equation 4, *d* denotes the average pairwise distances between the instance *x* and all other instances in the same cluster. *d*^*′*^ denotes the average pairwise distances between the instance *x* and all other instances in the nearest different cluster.

In order to evaluate the performance of population centroids estimation, we use three predictive performance metrics, i.e. MC-MCC, MC-F1 and ACC. Well-estimated centroids of cell groups should reflect the entire cell population’s properties – the corresponding assigned cell groups for testing instances should also bear high correctness. As shown in Equation 5, the Multi-Class Matthews Correlation Coefficient (MC-MCC) is an extension of the conventional MCC that was originally used to evaluate models’ performance in predicting labels of two classes, where *N* is the total number of instances, and 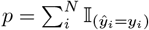 is the number of instances correctly predicted. *ŷ*_*i*_ is the predicted class of the *i*^*th*^ instance, and *y*_*i*_ is the true class. 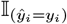 is an indicator function returning the value of 1 if *ŷ*_*i*_ = *y*_*i*_ and the value of 0 otherwise. 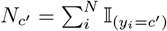 is the number of times class *c*′ occurred, and 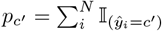 is the number of times class *c*′ is predicted. Analogously, the Multi-Class F1 score (MC-F1) is also an extension of the conventional F1 score into the multi-class settings, as shown in Equation 6, the subscript 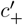 denotes the positive class, *C*′ is the set of classes, and *k*′ is the number of classes. Finally, the formula of accuracy (ACC) is shown in Equation 7. As all evaluation metrics are upper bounded by the value of 1 except the Calinski and Harabasz score, for consistency the values of Calinski and Harabasz score are scaled using the well-known MinMax scaling method so that all values lie in the interval [0, 1].

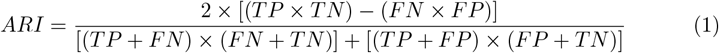

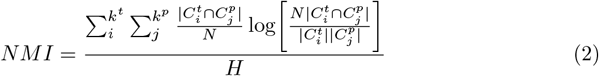

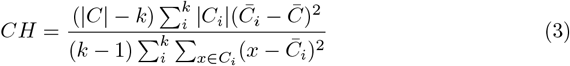

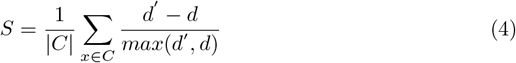

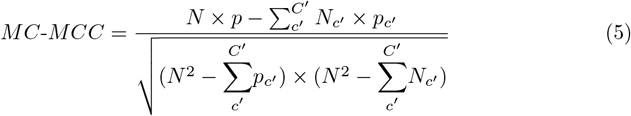

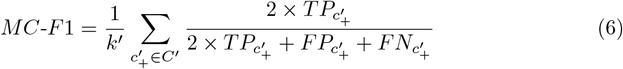

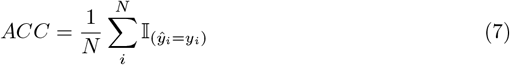

## Computational Experiments

### Evaluated scRNA-Seq contrastive learning methods

In this work, as shown in Table 1, we evaluate the generative encoder networks learned by five different types of state-of-the-art scRNA-Seq contrastive learning methods, i.e.

**Table 1.** The characteristics of scRNA-Seq contrastive learning algorithms evaluated in this work.

| Names | View Creation Methods | Loss Functions | References |
| --- | --- | --- | --- |
| Sup-GsRCL | Adding Gaussian noises | $\mathcal{L}^{SL} = \frac{1}{2N} \sum_{a=1}^{2N} \frac{-1}{\vert P\vert } \sum_{c \in P} \log \frac{\exp(\text{sim}(z_a, z_c)/\tau)}{\sum_{v=1}^{2N} (v \neq a) \exp(\text{sim}(z_a, z_v)/\tau)} (8)$ | [1] |
| Sup-RGMRCL | Masking random genes |  | [1] |
| Self-GsRCL | Adding Gaussian noises | $\mathcal{L}^{SSL} = \frac{-1}{2N} \sum_{a=1}^{2N} \log \frac{\exp(\text{sim}(z_a, z_b)/\tau)}{\sum_{v=1}^{2N} (v \neq a) \exp(\text{sim}(z_a, z_v)/\tau)} (9)$ | [1] |
| Self-RGMRCL | Masking random genes |  | [20] |
| AF-RCL | Using original samples as views | $\mathcal{L}_{AF}^{SL} = \frac{1}{N} \sum_{a=1}^N \frac{-1}{\vert H_a^+\vert } \sum_{h_q \in H_a^+} \log \frac{\exp(\text{sim}(h_a, h_q)/\tau)}{\exp(\text{sim}(h_a, h_q)/\tau) + \sum_{h_l \in H_a^-} \exp(\text{sim}(h_a, h_l)/\tau)} (10)$ | [22] |

\* $N$ denotes the number of training instances and *sim* denotes cosine similarity.

Supervised Gaussian noise augmentation-based scRNA-Seq Contrastive Learning (Sup-GsRCL), Supervised Random Gene Masking augmentation-based scRNA-Seq Contrastive Learning (Sup-RGMRCL), Self-supervised Gaussian noise augmentation-based scRNA-Seq Contrastive Learning (Self-GsRCL), Self-supervised Random Gene Masking augmentation-based scRNA-Seq Contrastive Learning (Self-RGMRCL), and Supervised Augmentation-Free single-cell RNA-Seq Contrastive Learning (AF-RCL). As introduced in [1], both Sup-GsRCL and Self-GsRCL methods create two views for each instance by respectively adding two different random Gaussian noise vectors to the original scRNA-Seq expression profiles. All Gaussian noise vectors are randomly drawn from a pre-defined Gaussian distribution *N* (0,1). Sup-GsRCL and Self-GsRCL then use the supervised and self-supervised contrastive learning loss functions to optimize network parameters, respectively shown in Equations 8 and 9. Both methods also showed good predictive performance when handling difficult cell-type identification tasks [1].

Analogously, both Sup-RGMRCL and Self-RGMRCL methods create two views for each instance by randomly masking *n* genes’ expression profiles. As introduced in [20], Self-RGMRCL showed state-of-the-art performance on scRNA-Seq clustering tasks. In terms of the discriminative model-based cell-type identification tasks, Self-RGMRCL with masking 3,000 random genes (i.e. Self-RGMRCL-3000) and Sup-RGMRCL with masking 5,000 random genes (i.e. Sup-RGMRCL-5000) both obtained good predictive accuracy when handling easy cell-type identification tasks [1]. The two methods also adopt the supervised and self-supervised contrastive learning loss functions to optimize network parameters, respectively.

In contrast to Sup-GsRCL, Sup-RGMRCL, Self-GsRCL and Self-RGMRCL, the AF-RCL method simplifies the conventional data augmentation-based view creating operation by adopting an augmentation-free approach – the original scRNA-Seq expression profiles are directly used as views to conduct contrastive learning using a different loss function (i.e. Equation 10). As introduced in [22], AF-RCL obtained the state-of-the-art predictive accuracy when handling different cell-type identification tasks in general.

### Experimental Datasets and Settings

18 different scRNA-Seq datasets [3, 24] are used to evaluate the performance of scRNA-Seq contrastive learning methods. As shown in Table 2, those 18 datasets that were obtained from different tissues (e.g. pancreas, lung and kidney) of different model organisms (e.g. Human and Mouse) consist of multiple cell-types. For example, the Segerstolpe (Human pancreas) dataset includes 13 different cell-types. Also, those 18 datasets include large numbers of genes but a relatively smaller number of cells. For example, the Xin (Human pancreas) dataset includes the expression profiles of the largest number of genes (i.e. 33,889), whilst the smallest number of genes included among those 18 datasets also reached 14,861 (i.e. the Baron Mouse pancreas dataset). However, almost all datasets include fewer than 4,000 cells, with the exception of one dataset, i.e. the PBMCBench 10Xv2 dataset that includes 6,444 cells. We split each individual dataset into two subsets using the 8:2 ratio, where 80% of the instances are used to conduct the well-known 5-fold cross validation, while the remaining 20% of the instances form the validation set for contrastive learning.

**Table 2.**
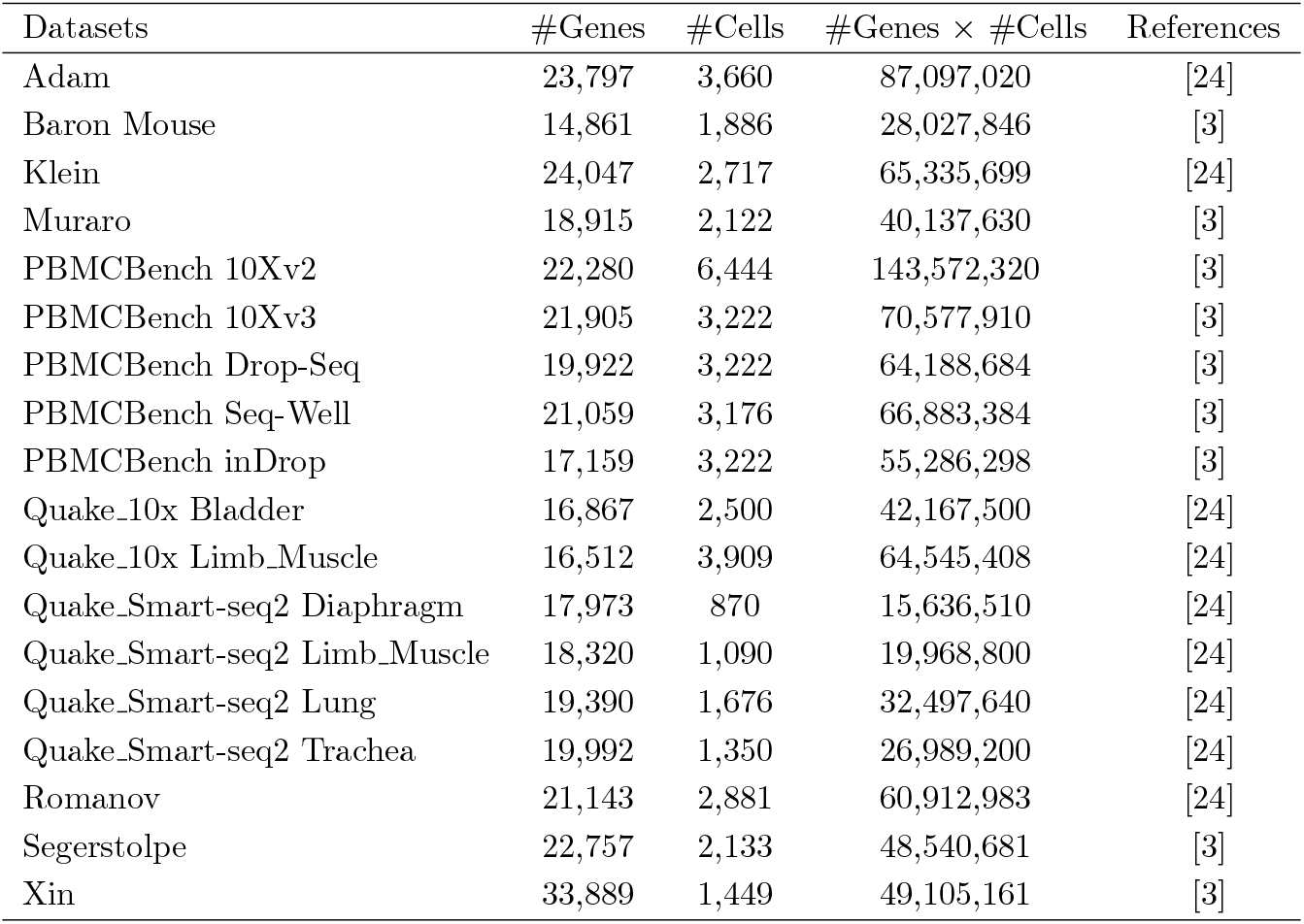
The characteristics of experimental scRNA-Seq datasets.

| Datasets | #Genes | #Cells | #Genes × #Cells | References |
| --- | --- | --- | --- | --- |
| Adam | 23,797 | 3,660 | 87,097,020 | [24] |
| Baron Mouse | 14,861 | 1,886 | 28,027,846 | [3] |
| Klein | 24,047 | 2,717 | 65,335,699 | [24] |
| Muraro | 18,915 | 2,122 | 40,137,630 | [3] |
| PBMCBench 10Xv2 | 22,280 | 6,444 | 143,572,320 | [3] |
| PBMCBench 10Xv3 | 21,905 | 3,222 | 70,577,910 | [3] |
| PBMCBench Drop-Seq | 19,922 | 3,222 | 64,188,684 | [3] |
| PBMCBench Seq-Well | 21,059 | 3,176 | 66,883,384 | [3] |
| PBMCBench inDrop | 17,159 | 3,222 | 55,286,298 | [3] |
| Quake_10x Bladder | 16,867 | 2,500 | 42,167,500 | [24] |
| Quake_10x Limb_Muscle | 16,512 | 3,909 | 64,545,408 | [24] |
| Quake_Smart-seq2 Diaphragm | 17,973 | 870 | 15,636,510 | [24] |
| Quake_Smart-seq2 Limb_Muscle | 18,320 | 1,090 | 19,968,800 | [24] |
| Quake_Smart-seq2 Lung | 19,390 | 1,676 | 32,497,640 | [24] |
| Quake_Smart-seq2 Trachea | 19,992 | 1,350 | 26,989,200 | [24] |
| Romanov | 21,143 | 2,881 | 60,912,983 | [24] |
| Segerstolpe | 22,757 | 2,133 | 48,540,681 | [3] |
| Xin | 33,889 | 1,449 | 49,105,161 | [3] |

### Experimental Results

Table 3 shows the ARI and NMI scores obtained by the scRNA-Seq embeddings generated by five different contrastive learning-based encoder generative networks and the original scRNA-Seq expression profiles. In general, when using the supervised contrastive learning paradigm, GsRCL (i.e. Sup-GsRCL) leads to better results than RGMRCL (i.e. Sup-RGMRCL-5000). Among those 18 datasets, the former obtains higher ARI and NMI scores in 12 and 8 datasets respectively, whilst the latter obtains better ARI and NMI scores in 6 and 8 datasets respectively. Both GsRCL and RGMRCL obtain the same NMI scores in 2 out of 18 datasets. GsRCL also leads to better results than AF-RCL, it obtains better ARI and NMI scores than AF-RCL in 12 and 10 datasets respectively, whilst AF-RCL obtains higher ARI and NMI scores in 4 and 6 datasets. Both GsRCL and AF-RCL obtain the same ARI and NMI scores in 2 and 1 datasets respectively. However, when using the self-supervised contrastive learning paradigm, RGMRCL (i.e. Self-RGMRCL-3000) leads to better results than GsRCL (i.e. Self-GsRCL). Among those 18 datasets, the former obtains higher ARI and NMI scores in almost all datasets, except two datasets where the latter obtains higher ARI scores. When comparing the cases using both types of contrastive learning paradigms (i.e. AF-RCL, Sup-GsRCL, Sup-RGMRCL-5000, Self-GsRCL and Self-RGMRCL-3000), GsRCL still leads to the overall best ARI scores, because Sup-GsRCL obtains the overall highest ARI values in 10 out of 18 datasets, whilst RGMRCL is the best-performing method in terms of NMI scores, since Sup-RGMRCL-5000 obtains the overall highest NMI values in 8 out of 18 datasets, as denoted by the underlines in Table 3. In addition, AF-RCL leads to better results than both Self-GsRCL and Self-RGMRCL-3000, whilst the original scRNA-Seq expression profiles obtain the overall worst results, compared with the embeddings generated by all different contrastive learning-based encoder networks.

**Table 3.** The Adjusted Rand Index (ARI) and Normalized Mutual Information (NMI) scores obtained by different scRNA-Seq contrastive learning generative encoder networks.

| Datasets | Selection Metrics | Supervised Contrastive Paradigm |  |  | Self-supervised Contrastive Paradigm |  | Raw Expression Profiles (Benchmark) |
| --- | --- | --- | --- | --- | --- | --- | --- |
|  |  | Augmentation free (AF-RCL) | Gaussian Noise (Sup-GsRCL) | Random 5,000 Genes Masking (Sup-RGMRCL-5000) | Gaussian Noise (Self-GsRCL) | Random 3,000 Genes Masking (Self-RGMRCL-3000) |  |
| Adam | ARI | 0.939 | 0.932 | **<u>0.944</u>** | 0.412 | **0.894** | 0.489 |
|  | NMI | 0.928 | 0.925 | **<u>0.936</u>** | 0.531 | **0.898** | 0.639 |
| Baron Mouse | ARI | 0.741 | **0.760** | 0.728 | 0.264 | **0.694** | 0.380 |
|  | NMI | 0.871 | **<u>0.874</u>** | 0.864 | 0.563 | **0.760** | 0.689 |
| Klein | ARI | **<u>0.994</u>** | **<u>0.994</u>** | 0.966 | 0.774 | **0.784** | 0.768 |
|  | NMI | **<u>0.992</u>** | 0.989 | 0.950 | 0.725 | **0.785** | 0.824 |
| Muraro | ARI | 0.943 | 0.954 | **<u>0.958</u>** | 0.467 | **0.912** | 0.685 |
|  | NMI | 0.923 | **<u>0.934</u>** | **<u>0.934</u>** | 0.645 | **0.873** | 0.814 |
| PBMCBench | ARI | **<u>0.655</u>** | 0.631 | 0.520 | 0.380 | **0.545** | 0.503 |
| 10Xv2 | NMI | **<u>0.760</u>** | 0.718 | 0.656 | 0.533 | **0.657** | 0.640 |
| PBMCBench | ARI | 0.651 | **<u>0.678</u>** | 0.580 | 0.510 | **0.585** | 0.549 |
| 10Xv3 | NMI | 0.775 | **<u>0.799</u>** | 0.709 | 0.626 | **0.702** | 0.705 |
| PBMCBench | ARI | **<u>0.653</u>** | 0.589 | 0.475 | 0.294 | **0.485** | 0.438 |
| Drop-Seq | NMI | **<u>0.701</u>** | 0.661 | 0.591 | 0.466 | **0.605** | 0.583 |
| PBMCBench | ARI | 0.000 | **<u>0.003</u>** | -0.002 | **-0.001** | -0.002 | -0.005 |
| Seq-Well | NMI | **<u>0.022</u>** | 0.020 | 0.021 | 0.021 | **<u>0.030</u>** | 0.024 |
| PBMCBench | ARI | -0.001 | **<u>0.003</u>** | 0.002 | **0.000** | -0.002 | 0.000 |
| inDrop | NMI | **<u>0.026</u>** | 0.021 | 0.024 | 0.021 | **0.023** | 0.023 |
| Quake_10x | ARI | 0.993 | 0.993 | **<u>0.997</u>** | 0.482 | **0.767** | 0.759 |
| Bladder | NMI | 0.983 | 0.985 | **<u>0.994</u>** | 0.571 | **0.823** | 0.815 |
| Quake_10x | ARI | 0.991 | **<u>0.994</u>** | 0.993 | 0.578 | **0.984** | 0.756 |
| Limb_Muscle | NMI | **<u>0.990</u>** | **<u>0.990</u>** | **<u>0.990</u>** | 0.663 | **0.977** | 0.856 |
| Quake_Smart-seq2 | ARI | 0.988 | **<u>0.997</u>** | 0.992 | 0.648 | **0.985** | 0.989 |
| Diaphragm | NMI | 0.985 | **<u>0.992</u>** | 0.984 | 0.789 | **0.984** | 0.985 |
| Quake_Smart-seq2 | ARI | 0.976 | 0.967 | **<u>0.983</u>** | 0.550 | **0.931** | 0.667 |
| Limb_Muscle | NMI | 0.964 | 0.959 | **<u>0.978</u>** | 0.703 | **0.900** | 0.829 |
| Quake_Smart-seq2 | ARI | 0.976 | 0.977 | **<u>0.982</u>** | 0.357 | **0.643** | 0.435 |
| Lung | NMI | 0.965 | 0.969 | **<u>0.972</u>** | 0.628 | **0.756** | 0.724 |
| Quake_Smart-seq2 | ARI | 0.979 | 0.986 | **<u>0.990</u>** | 0.512 | **0.676** | 0.578 |
| Trachea | NMI | 0.963 | 0.971 | **<u>0.978</u>** | 0.500 | **0.681** | 0.700 |
| Romanov | ARI | 0.854 | **<u>0.870</u>** | 0.866 | 0.342 | **0.664** | 0.494 |
|  | NMI | 0.814 | 0.824 | **<u>0.826</u>** | 0.448 | **0.716** | 0.545 |
| Segerstolpe | ARI | 0.871 | **<u>0.974</u>** | 0.805 | 0.220 | **0.650** | 0.484 |
|  | NMI | 0.906 | **<u>0.964</u>** | 0.911 | 0.463 | **0.770** | 0.726 |
| Xin | ARI | 0.987 | **<u>0.989</u>** | 0.982 | 0.560 | **0.739** | 0.627 |
|  | NMI | 0.964 | **<u>0.969</u>** | 0.964 | 0.620 | **0.746** | 0.766 |

**Bold text:** the highest value between the methods using the same contrastive learning paradigm.
Underline: the overall highest value among all six methods.

Table 4 shows the Silhouette scores obtained by different contrastive learning-based encoder generative networks according to ARI and NMI scores that are respectively used as the model selection metric. In general, when working with supervised contrastive learning paradigm, GsRCL (i.e. Sup-GsRCL) obtains higher Silhouette scores than RGMRCL (i.e. Sup-RGMRCL-5000) in 11 and 9 datasets according to ARI and NMI model selection metrics respectively. The latter obtains higher Silhouette scores in the rest of the datasets. Analogously, GsRCL obtains higher Silhouette scores than AF-RCL in 13 and 14 datasets respectively. When using the self-supervised contrastive learning paradigm, RGMRCL (i.e. Self-RGMRCL-3000) obtains higher Silhouette scores in almost all datasets, whilst GsRCL (i.e. Self-GsRCL) only obtains higher Silhouette scores in 2 datasets. In addition, Sup-GsRCL obtains the overall highest Silhouette scores in 7 and 5 datasets respectively using ARI and NMI as the model selection metric, whilst Sup-RGMRCL-5000 obtains the overall highest scores in 6 and 9 datasets. AF-RCL also leads to better Silhouette scores than both Self-GsRCL and Self-RGMRCL-3000, whilst the original scRNA-Seq expression profiles still obtain the overall worst results, compared with the embeddings generated by all different contrastive learning-based generative encoder networks.

**Table 4.** The Silhouette scores obtained by different scRNA-Seq contrastive learning generative encoder networks.

| Datasets | Selection Metrics | Supervised Contrastive Paradigm |  |  | Self-supervised Contrastive Paradigm |  | Raw Expression Profiles (Benchmark) |
| --- | --- | --- | --- | --- | --- | --- | --- |
|  |  | Augmentation free (AF-RCL) | Gaussian Noise (Sup-GsRCL) | Random 5,000 Genes Masking (Sup-RGMRCL-5000) | Gaussian Noise (Self-GsRCL) | Random 3,000 Genes Masking (Self-RGMRCL-3000) |  |
| Adam | ARI | 0.664 | 0.711 | **<u>0.732</u>** | 0.153 | **0.313** | 0.007 |
|  | NMI | 0.674 | 0.710 | **<u>0.732</u>** | 0.165 | **0.313** |  |
| Baron Mouse | ARI | **<u>0.481</u>** | 0.462 | 0.443 | 0.188 | **0.274** | -0.028 |
|  | NMI | 0.441 | 0.452 | **<u>0.454</u>** | 0.216 | **0.304** |  |
| Klein | ARI | 0.846 | **<u>0.849</u>** | 0.847 | 0.217 | **0.538** | 0.071 |
|  | NMI | 0.839 | **<u>0.858</u>** | 0.847 | 0.206 | **0.534** |  |
| Muraro | ARI | 0.721 | 0.799 | **<u>0.860</u>** | 0.142 | **0.462** | 0.005 |
|  | NMI | 0.721 | 0.803 | **<u>0.853</u>** | 0.152 | **0.479** |  |
| PBMCBench 10Xv2 | ARI | **<u>0.582</u>** | 0.565 | 0.348 | 0.271 | **0.287** | 0.005 |
|  | NMI | **<u>0.598</u>** | 0.576 | 0.387 | 0.253 | **0.296** |  |
| PBMCBench 10Xv3 | ARI | 0.701 | **<u>0.705</u>** | 0.610 | 0.278 | **0.379** | 0.049 |
|  | NMI | 0.677 | **<u>0.744</u>** | 0.582 | 0.274 | **0.401** |  |
| PBMCBench Drop-Seq | ARI | **<u>0.573</u>** | 0.442 | 0.409 | 0.219 | **0.231** | -0.058 |
|  | NMI | **<u>0.575</u>** | 0.453 | 0.407 | 0.247 | **0.265** |  |
| PBMCBench Seq-Well | ARI | **0.196** | 0.152 | 0.192 | **<u>0.229</u>** | 0.113 | -0.080 |
|  | NMI | 0.123 | 0.195 | **<u>0.284</u>** | **0.246** | 0.241 |  |
| PBMCBench inDrop | ARI | 0.190 | **0.269** | 0.191 | **<u>0.272</u>** | 0.153 | -0.026 |
|  | NMI | 0.184 | **0.279** | 0.230 | **<u>0.302</u>** | 0.143 |  |
| Quake_10x Bladder | ARI | 0.828 | **<u>0.909</u>** | 0.905 | 0.395 | **0.588** | 0.086 |
|  | NMI | 0.828 | 0.907 | **<u>0.936</u>** | 0.395 | **0.575** |  |
| Quake_10x Limb_Muscle | ARI | 0.850 | **<u>0.851</u>** | 0.849 | 0.295 | **0.432** | 0.050 |
|  | NMI | **<u>0.856</u>** | 0.852 | 0.850 | 0.285 | **0.432** |  |
| Quake_Smart-seq2 Diaphragm | ARI | 0.839 | 0.850 | **<u>0.870</u>** | 0.193 | **0.692** | 0.023 |
|  | NMI | 0.843 | 0.850 | **<u>0.863</u>** | 0.189 | **0.717** |  |
| Quake_Smart-seq2 Limb_Muscle | ARI | 0.826 | 0.803 | **<u>0.847</u>** | 0.159 | **0.614** | 0.017 |
|  | NMI | 0.826 | 0.803 | **<u>0.846</u>** | 0.256 | **0.583** |  |
| Quake_Smart-seq2 Lung | ARI | 0.791 | 0.794 | **<u>0.808</u>** | 0.165 | **0.274** | -0.014 |
|  | NMI | 0.762 | 0.794 | **<u>0.808</u>** | 0.176 | **0.513** |  |
| Quake_Smart-seq2 Trachea | ARI | 0.862 | **<u>0.878</u>** | 0.849 | 0.206 | **0.295** | 0.035 |
|  | NMI | 0.862 | **<u>0.878</u>** | 0.816 | 0.193 | **0.314** |  |
| Romanov | ARI | 0.616 | 0.662 | **<u>0.698</u>** | 0.164 | **0.434** | 0.004 |
|  | NMI | 0.634 | 0.659 | **<u>0.699</u>** | 0.139 | **0.438** |  |
| Segerstolpe | ARI | 0.518 | **<u>0.741</u>** | 0.576 | 0.119 | **0.157** | -0.046 |
|  | NMI | 0.554 | **<u>0.741</u>** | 0.579 | 0.115 | **0.170** |  |
| Xin | ARI | 0.799 | **<u>0.859</u>** | 0.749 | 0.202 | **0.213** | 0.001 |
|  | NMI | 0.819 | **<u>0.859</u>** | 0.749 | 0.183 | **0.251** |  |

**Bold text:** the highest value between the methods using the same contrastive learning paradigm.
Underline: the overall highest value among all six methods.

Table 5 shows the scaled Calinski and Harabasz scores obtained by different contrastive learning-based generative encoder networks according to two different model selection metrics (i.e. ARI and NMI scores). In general, when working with the supervised contrastive learning paradigm, GsRCL (i.e. Sup-GsRCL) outperforms RGMRCL (Sup-RGMRCL-5000) in 8 and 10 datasets respectively according to ARI and NMI model selection metrics. The latter obtains higher scores in 9 and 8 datasets. GsRCL still outperforms AF-RCL in 14 datasets according to both model selection metrics. When using the self-supervised contrastive learning paradigm, Self-RGMRCL-3000 still outperforms Self-GsRCL in almost all datasets, except 4 datasets where the latter obtained higher scores according to the ARI-based model selection metric. The latter also only obtains higher scores than the former in 2 datasets according to the NMI-based model selection metric. In addition, according to the ARI-based model selection metric, both Sup-GsRCL and Sup-RGMRCL-5000 obtain the overall highest scaled Calinski and Harabasz scores in 7 datasets. According to the NMI-based model selection metric, Sup-GsRCL obtains the overall highest scores in 8 datasets, whilst Sup-RGMRCL-5000 obtains the overall highest scores in 7 datasets.

**Table 5.** The scaled Calinski Harabasz scores obtained by different scRNA-Seq contrastive learning generative encoder networks.

| Datasets | Selection Metrics | Supervised Contrastive Paradigm |  |  | Self-supervised Contrastive Paradigm |  | Raw Expression Profiles (Benchmark) |
| --- | --- | --- | --- | --- | --- | --- | --- |
|  |  | Augmentation free (AF-RCL) | Gaussian Noise (Sup-GsRCL) | Random 5,000 Genes Masking (Sup-RGMRCL-5000) | Gaussian Noise (Self-GsRCL) | Random 3,000 Genes Masking (Self-RGMRCL-3000) |  |
| Adam | ARI | 0.070 | 0.120 | **<u>0.135</u>** | 0.006 | **0.017** | 0.003 |
|  | NMI | 0.074 | 0.121 | **<u>0.135</u>** | 0.007 | **0.017** |  |
| Baron Mouse | ARI | 0.062 | 0.048 | **<u>0.063</u>** | 0.003 | **0.010** | 0.000 |
|  | NMI | 0.063 | 0.053 | **<u>0.064</u>** | 0.005 | **0.011** |  |
| Klein | ARI | **<u>0.585</u>** | 0.462 | 0.484 | 0.010 | **0.058** | 0.007 |
|  | NMI | **<u>0.502</u>** | 0.491 | 0.484 | 0.010 | **0.054** |  |
| Muraro | ARI | 0.120 | 0.160 | **<u>0.264</u>** | 0.003 | **0.028** | 0.001 |
|  | NMI | 0.120 | 0.162 | **<u>0.241</u>** | 0.003 | **0.037** |  |
| PBMCBench | ARI | 0.157 | **<u>0.272</u>** | 0.104 | 0.020 | **0.077** | 0.004 |
| 10Xv2 | NMI | 0.172 | **<u>0.285</u>** | 0.142 | 0.018 | **0.080** |  |
| PBMCBench | ARI | 0.138 | **<u>0.197</u>** | 0.192 | 0.011 | **0.036** | 0.003 |
| 10Xv3 | NMI | 0.135 | **<u>0.194</u>** | 0.172 | 0.010 | **0.038** |  |
| PBMCBench | ARI | 0.059 | **<u>0.068</u>** | 0.055 | 0.012 | **0.022** | 0.000 |
| Drop-Seq | NMI | 0.063 | **<u>0.073</u>** | 0.055 | 0.015 | **0.025** |  |
| PBMCBench | ARI | 0.024 | **<u>0.025</u>** | 0.009 | **0.018** | 0.013 | 0.001 |
| Seq-Well | NMI | 0.018 | **<u>0.031</u>** | 0.011 | **0.015** | 0.014 |  |
| PBMCBench | ARI | 0.023 | **0.046** | 0.010 | **<u>0.084</u>** | 0.015 | 0.002 |
| inDrop | NMI | 0.024 | **<u>0.064</u>** | 0.010 | **0.032** | 0.016 |  |
| Quake_10x | ARI | 0.390 | 0.750 | **<u>0.896</u>** | 0.037 | **0.049** | 0.006 |
| Bladder | NMI | 0.390 | 0.935 | **<u>1.000</u>** | 0.037 | **0.046** |  |
| Quake_10x | ARI | 0.433 | **<u>0.512</u>** | 0.477 | 0.024 | **0.026** | 0.004 |
| Limb_Muscle | NMI | 0.482 | **<u>0.529</u>** | 0.488 | 0.024 | **0.026** |  |
| Quake_Smart-seq2 | ARI | 0.093 | **<u>0.143</u>** | 0.131 | 0.004 | **0.036** | 0.000 |
| Diaphragm | NMI | 0.106 | **<u>0.143</u>** | 0.136 | 0.005 | **0.041** |  |
| Quake_Smart-seq2 | ARI | **<u>0.123</u>** | 0.077 | 0.103 | 0.005 | **0.037** | 0.000 |
| Limb_Muscle | NMI | **<u>0.123</u>** | 0.077 | 0.102 | 0.004 | **0.036** |  |
| Quake_Smart-seq2 | ARI | 0.081 | 0.082 | **<u>0.086</u>** | 0.004 | **0.016** | 0.000 |
| Lung | NMI | 0.075 | 0.082 | **<u>0.086</u>** | 0.004 | **0.021** |  |
| Quake_Smart-seq2 | ARI | **<u>0.207</u>** | 0.173 | 0.173 | **0.010** | 0.006 | 0.001 |
| Trachea | NMI | **<u>0.207</u>** | 0.173 | 0.172 | 0.006 | **0.007** |  |
| Romanov | ARI | 0.057 | 0.075 | **<u>0.079</u>** | 0.006 | **0.027** | 0.002 |
|  | NMI | 0.061 | 0.074 | **<u>0.079</u>** | 0.004 | **0.028** |  |
| Segerstolpe | ARI | 0.076 | 0.104 | **<u>0.134</u>** | 0.003 | **0.004** | 0.000 |
|  | NMI | 0.078 | 0.104 | **<u>0.136</u>** | 0.002 | **0.005** |  |
| Xin | ARI | 0.228 | **<u>0.389</u>** | 0.134 | **0.008** | 0.005 | 0.000 |
|  | NMI | 0.310 | **<u>0.373</u>** | 0.134 | 0.008 | **0.010** |  |

**Bold text:** the highest value between the methods using the same contrastive learning paradigm.
Underline: the overall highest value among all six methods.

AF-RCL still leads to better scaled Calinski and Harabasz scores than both Self-GsRCL and Self-RGMRCL-3000 according to both model selection metrics. The original scRNA-Seq expression profiles still obtain the overall worst results.

In terms of the performance of estimating the centroids of different cell groups, as shown in Tables 6, 7, and 8, Sup-GsRCL is still the best-performing method, whilst both AF-RCL and Sup-RGMRCL-5000 also show competitive performance. When using the supervised contrastive learning paradigm, GsRCL (i.e. Sup-GsRCL) obtains higher MC-MCC values than RGMRCL (i.e. Sup-RGMRCL-5000) in 11 and 9 datasets, respectively using ARI and NMI scores as the model selection metric, whilst Sup-RGMRCL-5000 performs better than Sup-GsRCL in 7 and 9 datasets respectively. GsRCL also obtains higher MC-MCC values than AF-RCL in 9 datasets, whilst AF-RCL obtains better MC-MCC values in 8 datasets, according to the ARI-based model selection metric. Both methods obtain the same MC-MCC value in 1 dataset. However, according to the NMI-based model selection metric, AF-RCL outperforms GsRCL in 10 datasets, whilst GsRCL obtains higher MC-MCC values in 8 datasets. When using the self-supervised contrastive learning paradigm, RGMRCL (i.e. Self-RGMRCL-3000) outperforms GsRCL (i.e. Self-GsRCL) in the vast majority of datasets. The latter only obtains higher MC-MCC values in 4 out of 18 datasets using ARI-based model selection metric. In addition, Sup-GsRCL obtains the overall highest MC-MCC values in 6 and 4 datasets respectively according to the ARI and NMI-based model selection metric, whilst Sup-RGMRCL-5000 obtains the overall highest MC-MCC values in 6 datasets according to both model selection metrics. AF-RCL also obtains the overall highest MC-MCC values in 4 and 6 datasets respectively. The original scRNA-Seq expression profiles obtain the overall worst results.

**Table 6.** The multi-class MCC values obtained by different scRNA-Seq contrastive learning generative encoder networks.

| Datasets | Selection Metrics | Supervised Contrastive Paradigm |  |  | Self-supervised Contrastive Paradigm |  | Raw Expression Profiles (Benchmark) |
| --- | --- | --- | --- | --- | --- | --- | --- |
|  |  | Augmentation free (AF-RCL) | Gaussian Noise (Sup-GsRCL) | Random 5,000 Genes Masking (Sup-RGMRCL-5000) | Gaussian Noise (Self-GsRCL) | Random 3,000 Genes Masking (Self-RGMRCL-3000) |  |
| Adam | ARI | 0.961 | 0.957 | **<u>0.963</u>** | 0.846 | **0.948** | 0.592 |
|  | NMI | **<u>0.965</u>** | 0.961 | 0.963 | 0.847 | **0.948** |  |
| Baron Mouse | ARI | 0.947 | **<u>0.952</u>** | 0.942 | 0.840 | **0.905** | 0.920 |
|  | NMI | 0.952 | **<u>0.954</u>** | 0.946 | 0.778 | **0.930** |  |
| Klein | ARI | **<u>0.996</u>** | 0.992 | 0.983 | 0.911 | **0.930** | 0.740 |
|  | NMI | **<u>0.992</u>** | 0.990 | 0.983 | 0.909 | **0.935** |  |
| Muraro | ARI | 0.957 | **<u>0.969</u>** | 0.967 | 0.922 | **0.943** | 0.719 |
|  | NMI | 0.957 | **<u>0.970</u>** | 0.967 | 0.917 | **0.941** |  |
| PbmcBench 10Xv2 | ARI | **<u>0.799</u>** | 0.729 | 0.664 | **0.588** | 0.586 | 0.568 |
|  | NMI | **<u>0.803</u>** | 0.701 | 0.655 | 0.586 | **0.597** |  |
| PbmcBench 10Xv3 | ARI | 0.812 | **<u>0.833</u>** | 0.790 | **0.764** | 0.747 | 0.756 |
|  | NMI | 0.817 | **<u>0.828</u>** | 0.766 | 0.769 | **0.771** |  |
| PbmcBench Drop-Seq | ARI | **<u>0.777</u>** | 0.632 | 0.625 | **0.607** | 0.603 | 0.566 |
|  | NMI | **<u>0.775</u>** | 0.641 | 0.633 | 0.496 | **0.590** |  |
| PbmcBench Seq-Well | ARI | **0.013** | 0.008 | -0.006 | 0.017 | **0.020** | 0.024 |
|  | NMI | **0.013** | 0.001 | 0.012 | 0.021 | **<u>0.025</u>** |  |
| PbmcBench inDrop | ARI | **<u>0.005</u>** | -0.012 | 0.003 | **-0.003** | -0.012 | -0.001 |
|  | NMI | -0.007 | -0.006 | **-0.002** | -0.020 | **-0.015** |  |
| Quake_10x Bladder | ARI | 0.982 | **<u>0.993</u>** | 0.990 | 0.684 | **0.988** | 0.958 |
|  | NMI | 0.982 | 0.994 | **<u>0.996</u>** | 0.684 | **0.989** |  |
| Quake_10x Limb_Muscle | ARI | 0.990 | 0.995 | **<u>0.996</u>** | 0.831 | **0.992** | 0.927 |
|  | NMI | 0.995 | 0.994 | **<u>0.997</u>** | 0.835 | **0.992** |  |
| Quake_Smart-seq2 Diaphragm | ARI | 0.989 | **<u>0.991</u>** | 0.987 | 0.878 | **0.987** | 0.980 |
|  | NMI | **<u>0.993</u>** | 0.991 | 0.989 | 0.866 | **0.989** |  |
| Quake_Smart-seq2 Limb_Muscle | ARI | 0.983 | 0.976 | **<u>0.995</u>** | 0.724 | **0.959** | 0.943 |
|  | NMI | 0.983 | 0.976 | **<u>0.993</u>** | 0.953 | **0.968** |  |
| Quake_Smart-seq2 Lung | ARI | 0.972 | 0.978 | **<u>0.979</u>** | 0.774 | **0.882** | 0.935 |
|  | NMI | 0.964 | 0.978 | **<u>0.979</u>** | 0.790 | **0.941** |  |
| Quake_Smart-seq2 Trachea | ARI | 0.982 | 0.977 | **<u>0.984</u>** | 0.593 | **0.835** | 0.838 |
|  | NMI | **<u>0.982</u>** | 0.977 | **<u>0.982</u>** | 0.747 | **0.950** |  |
| Romanov | ARI | 0.885 | **<u>0.915</u>** | 0.903 | 0.744 | **0.875** | 0.650 |
|  | NMI | 0.889 | **<u>0.915</u>** | 0.905 | 0.806 | **0.872** |  |
| Segerstolpe | ARI | 0.942 | 0.959 | **<u>0.972</u>** | 0.748 | **0.864** | 0.809 |
|  | NMI | 0.948 | 0.959 | **<u>0.971</u>** | 0.794 | **0.875** |  |
| Xin | ARI | **0.967** | **0.967** | 0.946 | 0.829 | **0.852** | <u>0.992</u> |
|  | NMI | **0.975** | 0.967 | 0.946 | 0.814 | **0.926** |  |

**Bold text:** the highest value between the methods using the same contrastive learning paradigm.
Underline: the overall highest value among all six methods.

**Table 7.** The multi-class F1 scores obtained by different scRNA-Seq contrastive learning generative encoder networks.

| Datasets | Selection Metrics | Supervised Contrastive Paradigm |  |  | Self-supervised Contrastive Paradigm |  | Raw Expression Profiles (Benchmark) |
| --- | --- | --- | --- | --- | --- | --- | --- |
|  |  | Augmentation free (AF-RCL) | Gaussian Noise (Sup-GsRCL) | Random 5,000 Genes Masking (Sup-RGMRCL-5000) | Gaussian Noise (Self-GsRCL) | Random 3,000 Genes Masking (Self-RGMRCL-3000) |  |
| Adam | ARI | 0.962 | 0.959 | **<u>0.965</u>** | 0.860 | **0.952** | 0.658 |
|  | NMI | **<u>0.966</u>** | 0.962 | 0.965 | 0.860 | **0.952** |  |
| Baron Mouse | ARI | 0.753 | 0.790 | **<u>0.868</u>** | 0.695 | **0.864** | 0.837 |
|  | NMI | 0.776 | 0.798 | **<u>0.881</u>** | 0.611 | **0.868** |  |
| Klein | ARI | **<u>0.995</u>** | 0.991 | 0.986 | 0.918 | **0.935** | 0.789 |
|  | NMI | **<u>0.991</u>** | 0.989 | 0.986 | 0.918 | **0.940** |  |
| Muraro | ARI | 0.906 | **<u>0.949</u>** | 0.929 | 0.824 | **0.876** | 0.719 |
|  | NMI | 0.906 | **<u>0.949</u>** | 0.929 | 0.814 | **0.884** |  |
| PbmcBench 10Xv2 | ARI | 0.753 | **<u>0.775</u>** | 0.724 | 0.595 | **0.663** | 0.600 |
|  | NMI | 0.744 | **<u>0.770</u>** | 0.736 | 0.591 | **0.675** |  |
| PbmcBench 10Xv3 | ARI | 0.814 | 0.816 | **<u>0.843</u>** | 0.758 | **0.801** | 0.777 |
|  | NMI | 0.813 | 0.796 | **<u>0.823</u>** | 0.761 | **0.816** |  |
| PbmcBench Drop-Seq | ARI | **<u>0.729</u>** | 0.696 | 0.675 | 0.623 | **0.673** | 0.609 |
|  | NMI | **<u>0.739</u>** | 0.710 | 0.674 | 0.505 | **0.703** |  |
| PbmcBench Seq-Well | ARI | **<u>0.124</u>** | 0.087 | 0.114 | 0.067 | **0.072** | 0.123 |
|  | NMI | **<u>0.135</u>** | 0.089 | 0.116 | **0.069** | 0.068 |  |
| PbmcBench inDrop | ARI | **<u>0.133</u>** | 0.064 | 0.104 | 0.047 | **0.053** | 0.104 |
|  | NMI | **<u>0.128</u>** | 0.058 | 0.102 | 0.040 | **0.054** |  |
| Quake_10x Bladder | ARI | 0.955 | **<u>0.984</u>** | 0.962 | 0.561 | **0.969** | 0.905 |
|  | NMI | 0.955 | 0.989 | **<u>0.992</u>** | 0.561 | **0.972** |  |
| Quake_10x Limb_Muscle | ARI | 0.989 | 0.994 | **<u>0.996</u>** | 0.822 | **0.993** | 0.936 |
|  | NMI | 0.994 | 0.993 | **<u>0.997</u>** | 0.828 | **0.993** |  |
| Quake_Smart-seq2 Diaphragm | ARI | 0.989 | **<u>0.993</u>** | 0.981 | 0.861 | **0.981** | 0.984 |
|  | NMI | 0.991 | **<u>0.993</u>** | 0.987 | 0.840 | **0.989** |  |
| Quake_Smart-seq2 Limb_Muscle | ARI | 0.981 | 0.974 | **<u>0.995</u>** | 0.721 | **0.929** | 0.946 |
|  | NMI | 0.981 | 0.974 | **<u>0.994</u>** | **0.949** | 0.946 |  |
| Quake_Smart-seq2 Lung | ARI | 0.964 | 0.975 | **<u>0.976</u>** | 0.680 | **0.823** | 0.902 |
|  | NMI | 0.953 | 0.975 | **<u>0.976</u>** | 0.702 | **0.911** |  |
| Quake_Smart-seq2 Trachea | ARI | 0.983 | 0.981 | **<u>0.988</u>** | 0.677 | **0.878** | 0.891 |
|  | NMI | 0.983 | 0.981 | **<u>0.987</u>** | 0.800 | **0.966** |  |
| Romanov | ARI | 0.850 | **<u>0.895</u>** | 0.878 | 0.711 | **0.861** | 0.624 |
|  | NMI | 0.856 | **<u>0.893</u>** | 0.883 | 0.784 | **0.861** |  |
| Segerstolpe | ARI | 0.676 | 0.689 | **<u>0.739</u>** | 0.543 | **0.678** | 0.685 |
|  | NMI | 0.675 | 0.689 | **<u>0.737</u>** | 0.539 | **0.699** |  |
| Xin | ARI | **0.920** | 0.898 | 0.894 | **0.747** | 0.710 | <u>0.979</u> |
|  | NMI | **0.937** | 0.896 | 0.894 | 0.725 | **0.817** |  |

**Bold text:** the highest value between the methods using the same contrastive learning paradigm.
Underline: the overall highest value among all six methods.

**Table 8.** The accuracy (ACC) values obtained by different scRNA-Seq contrastive learning generative encoder networks.

| Datasets | Selection Metrics | Supervised Contrastive Paradigm |  |  | Self-supervised Contrastive Paradigm |  | Raw Expression Profiles (Benchmark) |
| --- | --- | --- | --- | --- | --- | --- | --- |
|  |  | Augmentation free (AF-RCL) | Gaussian Noise (Sup-GsRCL) | Random 5,000 Genes Masking (Sup-RGMRCL-5000) | Gaussian Noise (Self-GsRCL) | Random 3,000 Genes Masking (Self-RGMRCL-3000) |  |
| Adam | ARI | 0.967 | 0.963 | **<u>0.968</u>** | 0.866 | **0.955** | 0.639 |
|  | NMI | **<u>0.970</u>** | 0.966 | 0.968 | 0.867 | **0.955** |  |
| Baron Mouse | ARI | 0.962 | **<u>0.965</u>** | 0.958 | 0.883 | **0.931** | 0.941 |
|  | NMI | 0.966 | **<u>0.967</u>** | 0.961 | 0.826 | **0.950** |  |
| Klein | ARI | **<u>0.997</u>** | 0.994 | 0.988 | 0.936 | **0.948** | 0.805 |
|  | NMI | **<u>0.994</u>** | 0.993 | 0.988 | 0.934 | **0.953** |  |
| Muraro | ARI | 0.966 | **<u>0.976</u>** | 0.974 | 0.939 | **0.956** | 0.755 |
|  | NMI | 0.966 | **<u>0.976</u>** | 0.974 | 0.935 | **0.954** |  |
| PbmcBench 10Xv2 | ARI | **<u>0.840</u>** | 0.775 | 0.719 | **0.626** | 0.603 | 0.634 |
|  | NMI | **<u>0.842</u>** | 0.736 | 0.699 | **0.619** | 0.611 |  |
| PbmcBench 10Xv3 | ARI | 0.851 | **<u>0.865</u>** | 0.834 | **0.810** | 0.792 | 0.804 |
|  | NMI | 0.854 | **<u>0.859</u>** | 0.812 | 0.814 | **0.817** |  |
| PbmcBench Drop-Seq | ARI | **<u>0.833</u>** | 0.690 | 0.702 | **0.677** | 0.673 | 0.661 |
|  | NMI | **<u>0.832</u>** | 0.698 | 0.707 | 0.568 | **0.656** |  |
| PbmcBench Seq-Well | ARI | 0.301 | 0.137 | **<u>0.342</u>** | 0.071 | **0.128** | 0.202 |
|  | NMI | 0.322 | 0.172 | **<u>0.344</u>** | 0.071 | **0.120** |  |
| PbmcBench inDrop | ARI | **<u>0.272</u>** | 0.091 | 0.208 | 0.065 | **0.066** | 0.194 |
|  | NMI | **<u>0.277</u>** | 0.081 | 0.216 | 0.044 | **0.076** |  |
| Quake_10x Bladder | ARI | 0.990 | **<u>0.996</u>** | 0.995 | 0.782 | **0.994** | 0.976 |
|  | NMI | 0.990 | 0.997 | **<u>0.998</u>** | 0.782 | **0.994** |  |
| Quake_10x Limb_Muscle | ARI | 0.993 | 0.996 | **<u>0.997</u>** | 0.868 | **0.994** | 0.944 |
|  | NMI | 0.996 | 0.995 | **<u>0.997</u>** | 0.871 | **0.994** |  |
| Quake_Smart-seq2 Diaphragm | ARI | 0.993 | **<u>0.994</u>** | 0.991 | 0.917 | **0.991** | 0.987 |
|  | NMI | **<u>0.996</u>** | 0.994 | 0.993 | 0.908 | **0.993** |  |
| Quake_Smart-seq2 Limb_Muscle | ARI | 0.989 | 0.984 | **<u>0.997</u>** | 0.795 | **0.972** | 0.961 |
|  | NMI | 0.989 | 0.984 | **<u>0.995</u>** | 0.968 | **0.978** |  |
| Quake_Smart-seq2 Lung | ARI | 0.979 | **<u>0.984</u>** | **<u>0.984</u>** | 0.820 | **0.910** | 0.951 |
|  | NMI | 0.973 | **<u>0.984</u>** | **<u>0.984</u>** | 0.834 | **0.955** |  |
| Quake_Smart-seq2 Trachea | ARI | 0.990 | 0.987 | **<u>0.991</u>** | 0.731 | **0.906** | 0.906 |
|  | NMI | **<u>0.990</u>** | 0.987 | **<u>0.990</u>** | 0.840 | **0.971** |  |
| Romanov | ARI | 0.914 | **<u>0.936</u>** | 0.927 | 0.803 | **0.905** | 0.728 |
|  | NMI | 0.917 | **<u>0.936</u>** | 0.929 | 0.854 | **0.903** |  |
| Segerstolpe | ARI | 0.955 | 0.968 | **<u>0.978</u>** | 0.791 | **0.888** | 0.841 |
|  | NMI | 0.960 | 0.968 | **<u>0.978</u>** | 0.832 | **0.896** |  |
| Xin | ARI | **0.982** | **0.982** | 0.971 | 0.900 | **0.920** | <u>0.996</u> |
|  | NMI | **0.986** | 0.982 | 0.971 | 0.890 | **0.960** |  |

**Bold text:** the highest value between the methods using the same contrastive learning paradigm.
Underline: the overall highest value among all six methods.

However, in terms of MC-F1 scores, Sup-RGMRCL-5000 outperforms Sup-GsRCL in 10 and 11 datasets, respectively using ARI and NMI scores as the model selection metric, whilst the latter obtains higher MC-F1 scores than the former in 8 and 7 datasets respectively. But Sup-GsRCL outperforms AF-RCL in 10 and 8 datasets according to the ARI-based and NMI-based model selection metric respectively.

Self-RGMRCL-3000 also outperforms Self-GsRCL in almost all datasets. Self-GsRCL only obtains higher MC-F1 scores in 1 and 2 datasets respectively according to ARI and NMI model selection metrics. Sup-RGMRCL-5000 also obtains the overall highest MC-F1 scores in 8 datasets according to both ARI and NMI-based model selection metrics, whilst Sup-GsRCL only obtains the overall highest MC-F1 scores in 5 and 4 datasets respectively. Analogously, AF-RCL obtains the overall highest MC-F1 scores in 4 and 5 datasets respectively. The original scRNA-Seq expression profiles obtain the overall worst results.

Finally, in terms of ACC values, when using the supervised contrastive learning paradigm, GsRCL (i.e. Sup-GsRCL), RGMRCL (i.e. Sup-RGMRCL-5000), and AF-RCL all show competitive performance. Sup-RGMRCL-5000 obtains higher ACC values than Sup-GsRCL in 8 and 9 datasets, respectively according to ARI and NMI model selection metrics, whilst the latter obtains higher ACC values in 9 and 8 datasets respectively. Both methods obtain the same ACC value in 1 out of 18 datasets.

AF-RCL also obtains higher ACC values than Sup-GsRCL in 8 and 11 datasets respectively, whilst the latter obtains better ACC values in 9 and 7 datasets respectively. Both methods obtain the same ACC value in 1 out of 18 datasets, according to the ARI-based model selection metric. When using the self-supervised contrastive learning paradigm, Self-RGMRCL-3000 still outperforms Self-GsRCL in the vast majority of the datasets. The latter only obtains higher ACC values in 3 datasets according to the ARI-based model selection metric. Sup-GsRCL obtains the overall highest ACC values in 7 and 5 datasets, whilst Sup-RGMRCL-5000 obtains the overall highest ACC values in 7 datasets according to both model selection metrics. AF-RCL also obtains the overall highest ACC values in 4 and 7 datasets respectively. The original scRNA-Seq expression profiles still obtain the overall worst results.

## Discussion

### Statistical Significance Tests on Computational Results

We first conduct statistical significance tests on the ARI, NMI, Silhouette and scaled Calinski Harabasz scores obtained by different methods. Table 9 shows the results of the Friedman tests with Holm’s *post-hoc* corrections. In general, Sup-GsRCL is the overall best-performing method that obtains the best average rankings and is used as the control method for all the above metrics. AF-RCL and Sup-RGMRCL-5000 also obtain the same best average rankings as Sup-GsRCL in terms of the NMI score and NMI value-selected Silhouette score respectively. When using Sup-GsRCL as the control method, it significantly outperforms Self-RGMRCL-3000, Self-GsRCL and the benchmark method according to all the above metrics, as denoted by the underlines. However, there is no significant difference between Sup-GsRCL and AF-RCL, or between Sup-GsRCL and Sup-RGMRCL-5000.

**Table 9.** The statistical testing results for comparisons of different scRNA-Seq contrastive learning generative encoder networks.

| Adjusted Rand Index (ARI) |  |  |  | Normalized Mutual Information (NMI) |  |  |  |
| --- | --- | --- | --- | --- | --- | --- | --- |
| Friedman test (p-value: 5.60E-13) |  |  |  | Friedman test (p-value: 1.73E-11) |  |  |  |
| Method | Ave. Rank | P-value | Adjusted $\alpha$ | Method | Ave. Rank | P-value | Adjusted $\alpha$ |
| Sup-GsRCL (ctrl.) | 1.611 | N/A | N/A | Sup-GsRCL (ctrl.) | 2.222 | N/A | N/A |
| Sup-RGMRCL-5000 | 2.361 | 2.29E-01 | 5.00E-02 | AF-RCL | 2.222 | 1.00E+00 | 5.00E-02 |
| AF-RCL | 2.444 | 1.81E-01 | 2.50E-02 | Sup-RGMRCL-5000 | 2.333 | 8.59E-01 | 2.50E-02 |
| Self-RGMRCL-3000 | 4.028 | <u>1.06E-04</u> | 1.67E-02 | Self-RGMRCL-3000 | 3.944 | <u>5.75E-03</u> | 1.67E-02 |
| Benchmark | 4.917 | <u>1.15E-07</u> | 1.25E-02 | Benchmark | 4.389 | <u>5.12E-04</u> | 1.25E-02 |
| Self-GsRCL | 5.639 | <u>1.06E-10</u> | 1.00E-02 | Self-GsRCL | 5.889 | <u>4.11E-09</u> | 1.00E-02 |

| Silhouette Score |  |  |  |  |  |  |  |
| --- | --- | --- | --- | --- | --- | --- | --- |
| ARI-based model selection |  |  |  | NMI-based model selection |  |  |  |
| Friedman test (p-value: 7.91E-14) |  |  |  | Friedman test (p-value: 7.91E-14) |  |  |  |
| Method | Ave. Rank | P-value | Adjusted $\alpha$ | Method | Ave. Rank | P-value | Adjusted $\alpha$ |
| Sup-GsRCL (ctrl.) | 1.778 | N/A | N/A | Sup-GsRCL (ctrl.) | 1.889 | N/A | N/A |
| Sup-RGMRCL-5000 | 2.167 | 5.33E-01 | 5.00E-02 | Sup-RGMRCL-5000 | 1.889 | 1.00E+00 | 5.00E-02 |
| AF-RCL | 2.389 | 3.27E-01 | 2.50E-02 | AF-RCL | 2.611 | 2.47E-01 | 2.50E-02 |
| Self-RGMRCL-3000 | 4.111 | <u>1.83E-04</u> | 1.67E-02 | Self-RGMRCL-3000 | 4.000 | <u>7.11E-04</u> | 1.67E-02 |
| Self-GsRCL | 4.556 | <u>8.41E-06</u> | 1.25E-02 | Self-GsRCL | 4.611 | <u>1.27E-05</u> | 1.25E-02 |
| Benchmark | 6.000 | <u>1.28E-11</u> | 1.00E-02 | Benchmark | 6.000 | <u>4.33E-11</u> | 1.00E-02 |

| Calinski Harabasz Score (scaled) |  |  |  |  |  |  |  |
| --- | --- | --- | --- | --- | --- | --- | --- |
| ARI-based model selection |  |  |  | NMI-based model selection |  |  |  |
| Friedman test (p-value: 8.67E-14) |  |  |  | Friedman test (p-value: 2.27E-14) |  |  |  |
| Method | Ave. Rank | P-value | Adjusted $\alpha$ | Method | Ave. Rank | P-value | Adjusted $\alpha$ |
| Sup-GsRCL (ctrl.) | 1.833 | N/A | N/A | Sup-GsRCL (ctrl.) | 1.667 | N/A | N/A |
| Sup-RGMRCL-5000 | 2.111 | 6.56E-01 | 5.00E-02 | Sup-RGMRCL-5000 | 2.222 | 3.73E-01 | 5.00E-02 |
| AF-RCL | 2.389 | 3.73E-01 | 2.50E-02 | AF-RCL | 2.389 | 2.47E-01 | 2.50E-02 |
| Self-RGMRCL-3000 | 4.111 | <u>2.60E-04</u> | 1.67E-02 | Self-RGMRCL-3000 | 4.000 | <u>1.83E-04</u> | 1.67E-02 |
| Self-GsRCL | 4.556 | <u>1.27E-05</u> | 1.25E-02 | Self-GsRCL | 4.722 | <u>9.59E-07</u> | 1.25E-02 |
| Benchmark | 6.000 | <u>2.36E-11</u> | 1.00E-02 | Benchmark | 6.000 | <u>3.68E-12</u> | 1.00E-02 |

| Multi-class MCC Value |  |  |  |  |  |  |  |
| --- | --- | --- | --- | --- | --- | --- | --- |
| ARI-based model selection |  |  |  | NMI-based model selection |  |  |  |
| Friedman test (p-value: 1.04E-08) |  |  |  | Friedman test (p-value: 3.32E-09) |  |  |  |
| Method | Ave. Rank | P-value | Adjusted $\alpha$ | Method | Ave. Rank | P-value | Adjusted $\alpha$ |
| Sup-GsRCL (ctrl.) | 2.222 | N/A | N/A | AF-RCL (ctrl.) | 2.250 | N/A | N/A |
| AF-RCL | 2.361 | 8.24E-01 | 5.00E-02 | Sup-GsRCL | 2.333 | 8.94E-01 | 5.00E-02 |
| Sup-RGMRCL-5000 | 2.361 | 8.24E-01 | 2.50E-02 | Sup-RGMRCL-5000 | 2.444 | 7.55E-01 | 2.50E-02 |
| Self-RGMRCL-3000 | 4.278 | <u>9.80E-04</u> | 1.67E-02 | Self-RGMRCL-3000 | 3.806 | <u>1.26E-02</u> | 1.67E-02 |
| Benchmark | 4.611 | <u>1.28E-04</u> | 1.25E-02 | Benchmark | 4.778 | <u>5.05E-05</u> | 1.25E-02 |
| Self-GsRCL | 5.167 | <u>2.34E-06</u> | 1.00E-02 | Self-GsRCL | 5.389 | <u>4.82E-07</u> | 1.00E-02 |

| Multi-class F1 Score |  |  |  |  |  |  |  |
| --- | --- | --- | --- | --- | --- | --- | --- |
| ARI-based model selection |  |  |  | NMI-based model selection |  |  |  |
| Friedman test (p-value: 6.87E-10) |  |  |  | Friedman test (p-value: 2.52E-09) |  |  |  |
| Method | Ave. Rank | P-value | Adjusted $\alpha$ | Method | Ave. Rank | P-value | Adjusted $\alpha$ |
| Sup-RGMRCL-5000 (ctrl.) | 2.111 | N/A | N/A | Sup-RGMRCL-5000 (ctrl.) | 2.111 | N/A | N/A |
| Sup-GsRCL | 2.278 | 7.89E-01 | 5.00E-02 | AF-RCL | 2.444 | 5.93E-01 | 5.00E-02 |
| AF-RCL | 2.556 | 4.76E-01 | 2.50E-02 | Sup-GsRCL | 2.583 | 4.49E-01 | 2.50E-02 |
| Self-RGMRCL-3000 | 4.139 | <u>1.15E-03</u> | 1.67E-02 | Self-RGMRCL-3000 | 3.722 | <u>9.78E-03</u> | 1.67E-02 |
| Benchmark | 4.250 | <u>6.04E-04</u> | 1.25E-02 | Benchmark | 4.528 | <u>1.06E-04</u> | 1.25E-02 |
| Self-GsRCL | 5.667 | <u>1.19E-08</u> | 1.00E-02 | Self-GsRCL | 5.611 | <u>1.99E-08</u> | 1.00E-02 |

| Accuracy |  |  |  |  |  |  |  |
| --- | --- | --- | --- | --- | --- | --- | --- |
| ARI-based model selection |  |  |  | NMI-based model selection |  |  |  |
| Friedman test (p-value: 7.91E-12) |  |  |  | Friedman test (p-value: 7.53E-12) |  |  |  |
| Method | Ave. Rank | P-value | Adjusted $\alpha$ | Method | Ave. Rank | P-value | Adjusted $\alpha$ |
| Sup-RGMRCL-5000 (ctrl.) | 2.056 | N/A | N/A | AF-RCL (ctrl.) | 1.972 | N/A | N/A |
| Sup-GsRCL | 2.111 | 9.29E-01 | 5.00E-02 | Sup-RGMRCL-5000 | 2.194 | 7.22E-01 | 5.00E-02 |
| AF-RCL | 2.250 | 7.55E-01 | 2.50E-02 | Sup-GsRCL | 2.306 | 5.93E-01 | 2.50E-02 |
| Self-RGMRCL-3000 | 4.444 | <u>1.28E-04</u> | 1.67E-02 | Self-RGMRCL-3000 | 4.194 | <u>3.66E-04</u> | 1.67E-02 |
| Benchmark | 4.639 | <u>3.43E-05</u> | 1.25E-02 | Benchmark | 4.778 | <u>6.83E-06</u> | 1.25E-02 |
| Self-GsRCL | 5.500 | <u>3.33E-08</u> | 1.00E-02 | Self-GsRCL | 5.556 | <u>9.13E-09</u> | 1.00E-02 |

Underline: the p-value that is lower than the corresponding significance level.

In terms of the contrastive learning models’ performance of estimating cell-type centroids, as also shown in Table 9, according to the MC-MCC values, Sup-GsRCL obtains the best average ranking when using the ARI-based model selection and shows significant differences to other methods except AF-RCL and Sup-RGMRCL-5000.

When using the NMI-based model selection metric, AF-RCL obtains the best average ranking and shows significantly higher MCC values than other methods except Sup-GsRCL and Sup-RGMRCL-5000. In terms of the MC-F1 scores (either using the ARI or NMI model selection metrics), Sup-RGMRCL-5000 obtains the best average rankings and shows significant differences to other methods except Sup-GsRCL and AF-RCL. Analogously, Sup-RGMRCL-5000 also shows significantly higher ACC values (according to the ARI-based model selection metric) than Self-RGMRCL-3000, Benchmark and Self-GsRCL. When using the NMI-based model selection metric,

AF-RCL obtains the best average ranking and shows significant differences to other methods except Sup-RGMRCL-5000 and Sup-GsRCL.

### Correlation coefficient between the performance of different models and the characteristics of scRNA-Seq datasets

We then further investigate the associations between the performance of contrastive learning generative encoder networks and the characteristics of scRNA-Seq datasets. As shown in Figure 2, in terms of the cell group estimation performance metrics,

**Fig 2.**
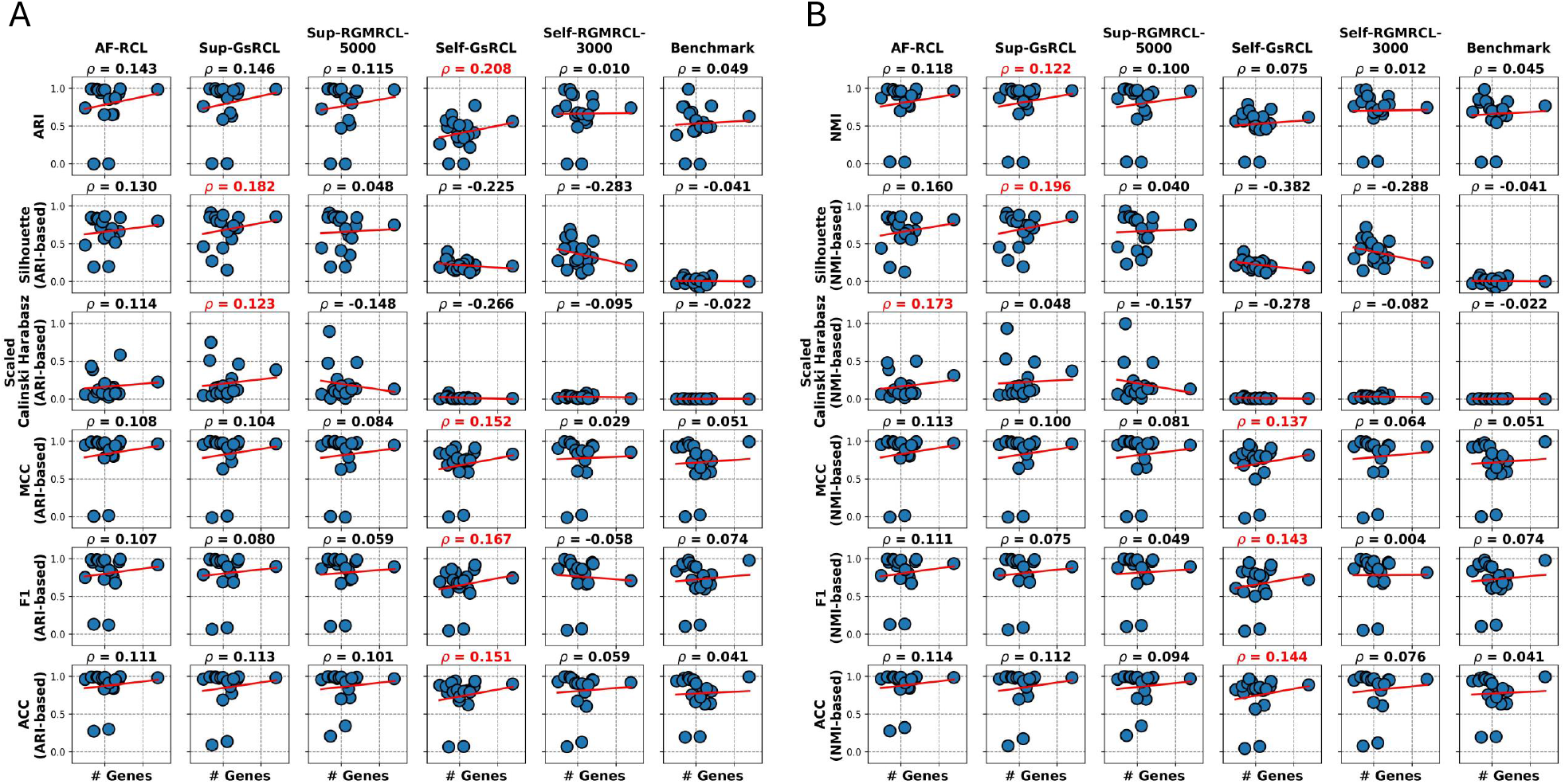
Scatter plots showing the correlations between (A) ARI-based performance and (B) NMI-based performance of different contrastive learning methods and the number of genes of the scRNA-Seq datasets. The highest Pearson correlation coefficient value is highlighted in red.

Sup-GsRCL shows the best Pearson correlation coefficient values between the number of genes and three metrics, i.e. 0.122 for NMI score, 0.182 and 0.196 for Silhouette score (using either ARI or NMI scores as the model selection metrics respectively), and 0.123 for the ARI value-selected Calinski Harabasz score. Sup-GsRCL also obtains the second-best correlation coefficient values, i.e. 0.146 for the ARI score and 0.048 for the NMI value-selected Calinski Harabasz score. All those correlation coefficient values are positive, suggesting that Sup-GsRCL has the strongest robustness against the high dimensionality issue of scRNA-Seq data. The second-best performing method is AF-RCL, which shows the highest positive correlation coefficient value (i.e. 0.173) for the NMI value-selected Calinski Harabasz score, and obtains the second-highest positive correlation coefficient values for all other metrics except the ARI score. Analogously, AF-RCL shows strong robustness against the high dimensionality issue of scRNA-Seq data. Self-GsRCL also shows the highest positive correlation coefficient value (i.e. 0.208) for the ARI score. However, in terms of the NMI score, it only obtains a very weak positive correlation coefficient value (i.e. 0.075). It also showed negative linear correlations for the other two metrics. Analogously, all other methods show either negative linear correlations or almost no linear correlations between their performance and the high dimensionality of scRNA-Seq data. In terms of those centroid estimation performance metrics (i.e. MC-MCC, MC-F1 and ACC), Self-GsRCL shows the best associations with the feature dimensionality, though the corresponding correlation coefficient values are not high, i.e. ranging from 0.137 to 0.167. All the other methods also have very weak correlation coefficient values.

In terms of the associations between the performance of contrastive learning methods and the number of cells (i.e. sample size) included in scRNA-Seq datasets, as shown in Figure 3, Self-GsRCL obtained the best correlation coefficient values for the ARI score (i.e. −0.240) and the Silhouette score with both types of model selection metrics (i.e. 0.400 and 0.282 for ARI and NMI respectively). Self-RGMRCL-3000 obtained the best correlation coefficient values for the NMI score (i.e. −0.250) and the Calinski Harabasz score with both types of model selection metrics (i.e. 0.507 and 0.501 for ARI and NMI respectively). Note that all methods obtained negative correlation coefficient values for ARI and NMI scores, suggesting the natural difficulty of distribution estimation tasks when the sample size becomes large. Analogously, the correlation coefficient values for the Silhouette score obtained by all methods except Self-GsRCL are also negative, though all methods obtained positive correlation coefficient values for the Calinski Harabasz score. In terms of the associations between cell group centroids estimation performance and the sample size, all methods also obtained negative correlation coefficient values, though AF-RCL shows the strongest robustness due to its highest correlation coefficient values for the NMI value-selected MC-MCC and the ACC scores with both types of model selection metrics. Self-GsRCL still shows strong robustness due to its highest correlation coefficient values for MC-MCC values and MC-F1 scores (the ones selected by ARI values). In addition, Sup-GsRCL obtains the best correlation coefficient value for the MC-F1 score using the NMI score as the model selection metric. Those correlation coefficient values are also all better than the ones obtained by the benchmark method, suggesting the usefulness of contrastive learning methods in general.

**Fig 3.**
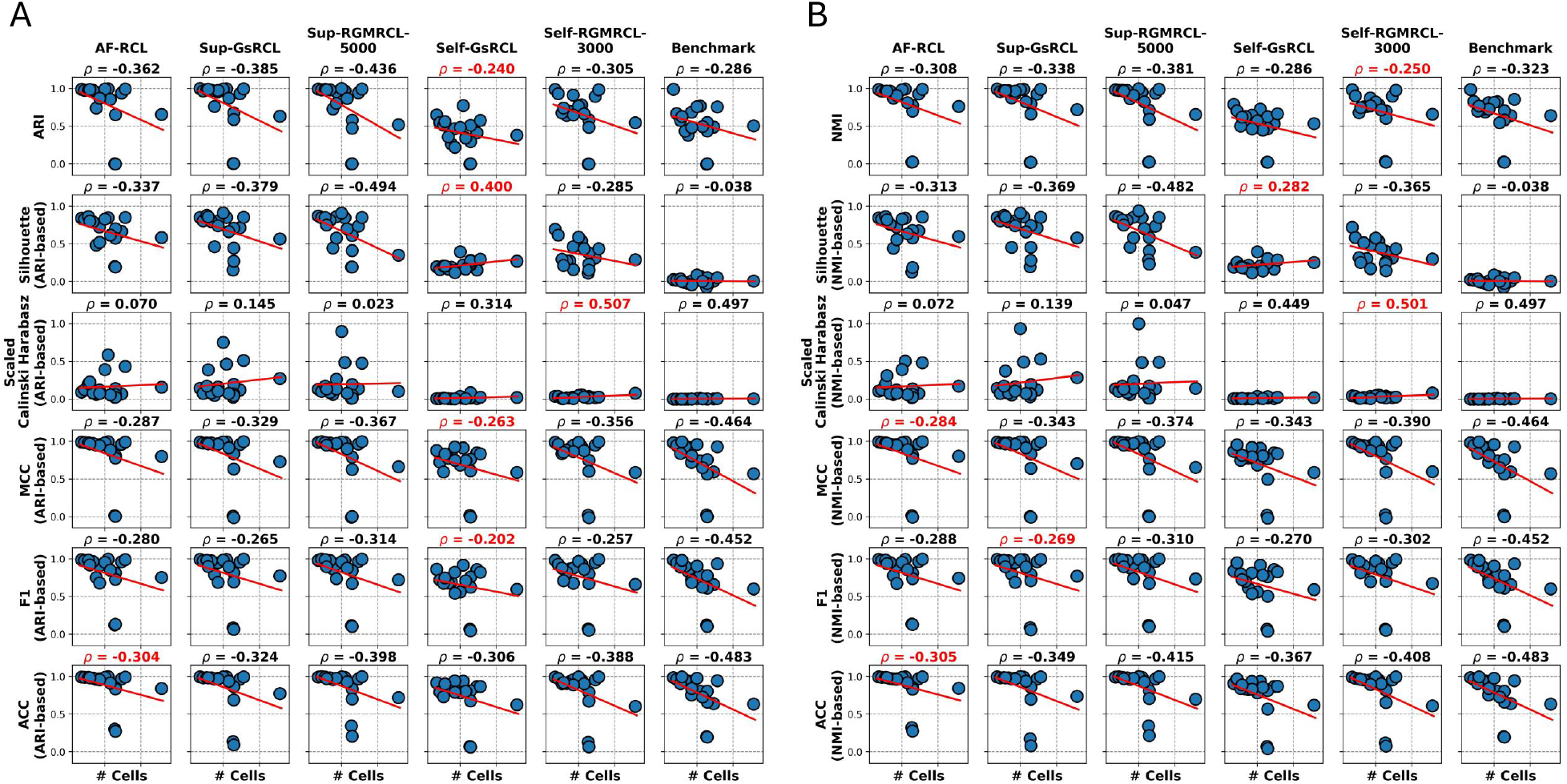
Scatter plots showing the correlations between (A) ARI-based performance and (B) NMI-based performance of different contrastive learning methods and the number of cells of the scRNA-Seq datasets. The highest Pearson correlation coefficient value is highlighted in red.

Those patterns discussed above are also observed when evaluating the correlation coefficient values between the full dimensionality (i.e. # Genes × # Cells) of datasets and different evaluation metrics. As shown in Figure 4, Self-GsRCL still obtains the highest correlation coefficient values for the ARI score (i.e. −0.152) and the Silhouette score with both selection metrics (i.e. 0.267 and 0.119 for ARI and NMI respectively). Self-RGMRCL-3000 also still obtains the best correlation coefficient values for the NMI score (i.e. −0.210) and the Calinski Harabasz score with both selection metrics (i.e. 0.497 and 0.496 for ARI and NMI respectively). In terms of the associations between the cell group centroids estimation performance and the full dimensionality of datasets, Self-GsRCL obtains the highest correlation coefficient values in terms of MC-MCC and ACC values (the ones selected by the ARI values), and the MC-F1 score with both model selection metrics. AF-RCL also obtains the highest correlation coefficient values for the MC-MCC and ACC values (the ones selected by the NMI values).

**Fig 4.**
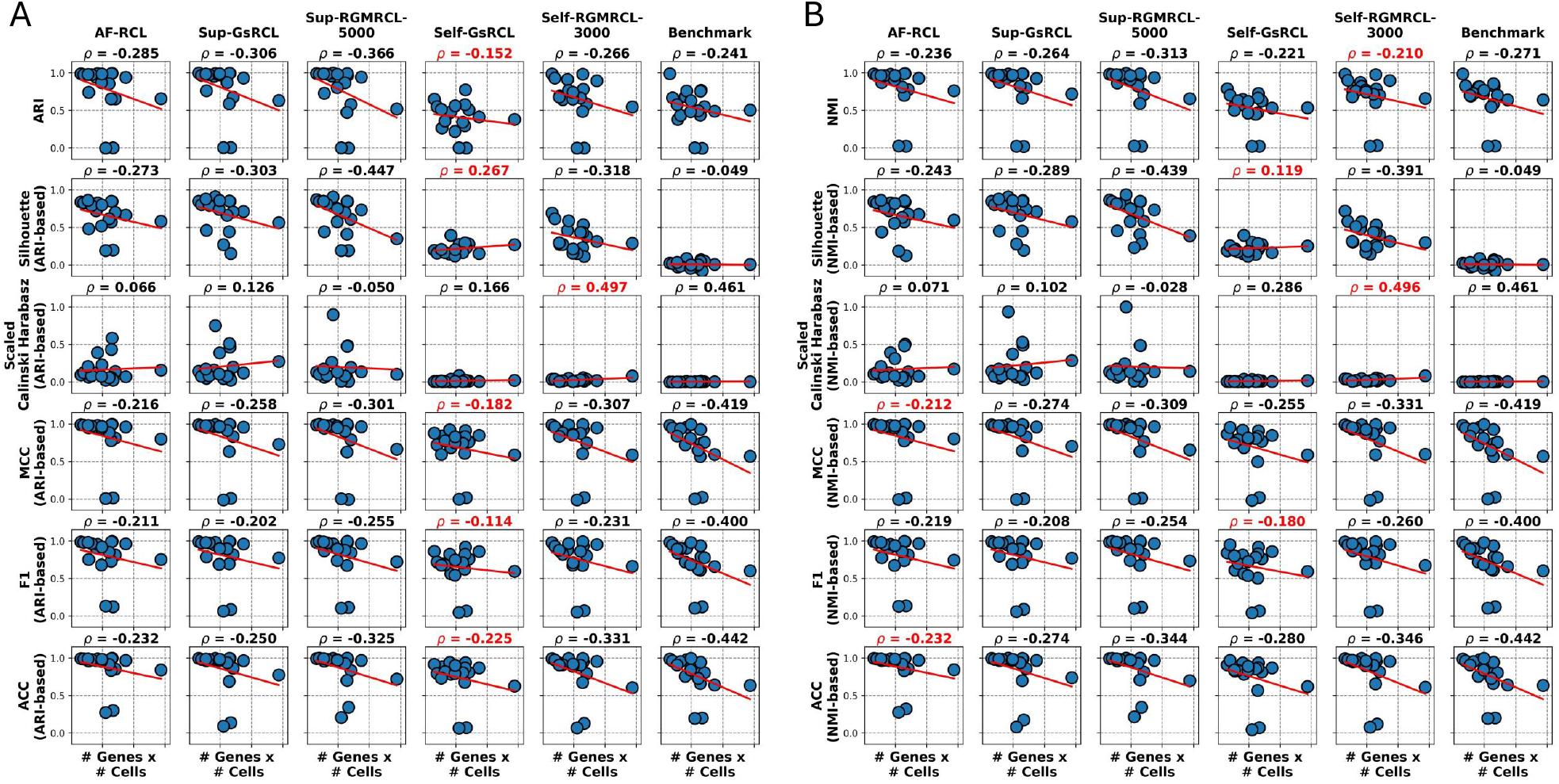
Scatter plots showing the correlations between (A) ARI-based performance and (B) NMI-based performance of different contrastive learning methods and the full dimensionality (i.e. # Genes × # Cells) of the scRNA-Seq datasets. The highest Pearson correlation coefficient value is highlighted in red.

## Conclusion

In conclusion, supervised scRNA-Seq contrastive learning methods lead to better performance than self-supervised scRNA-Seq contrastive learning methods in estimating the intrinsic distributions of different cell-types, whilst Sup-GsRCL is the overall best-performing method. In addition, among the self-supervised scRNA-Seq contrastive learning methods discussed in this work, Self-RGMRCL-3000 performs better than Self-GsRCL. Future research directions could focus on developing new supervised contrastive learning methods that exploit more powerful data augmentation strategies to further improve the performance of generative encoder networks on intrinsic cell-types distribution estimation.

## Acknowledgements

The authors acknowledge the support of the School of Computing and Mathematical Sciences and the Birkbeck GTA programme.

## Data availability

The single-cell RNA-seq datasets used in this work can be downloaded at https://doi.org/10.5281/zenodo.8087611 and the pre-trained encoders can be downloaded at https://doi.org/10.5281/zenodo.16955668 and at https://doi.org/10.5281/zenodo.16956501.

## Code availability

The source code can be downloaded at https://github.com/ibrahimsaggaf/scRCL-G.

## References

1. Alsaggaf I, Buchan D, Wan C. Improving cell type identification with Gaussian noise-augmented single-cell RNA-seq contrastive learning. Briefings in Functional Genomics. 2024;23:441–451.

2. Yu L, Cao Y, Yang JYH, Yang P. Benchmarking clustering algorithms on estimating the number of cell types from single-cell RNA-sequencing data. Genome Biology. 2022;23.

3. Abdelaal T, Michielsen L, Cats D, Hoogduin D, Mei H, Reinders MJT, et al. A comparison of automatic cell identification methods for single-cell RNA sequencing data. Genome Biology. 2019;20.

4. Yuan Q, Duren Z. Inferring gene regulatory networks from single-cell multiome data using atlas-scale external data. Nature Biotechnology. 2024;.

5. Keyl P, Bischoff P, Dernbach G, Bockmayr M, Fritz R, Horst D, et al. Single-cell gene regulatory network prediction by explainable AI. Nucleic Acids Research. 2023;51:e20.

6. Mao G, Pang Z, Zuo K, Wang Q, Pei X, Chen X, et al. Predicting gene regulatory links from single-cell RNA-seq data using graph neural networks. Briefings in Bioinformatics. 2023;24:bbad414.

7. de Sande BV, Lee JS, Mutasa-Gottgens E, Naughton B, Bacon W, Manning J, et al. Applications of single-cell RNA sequencing in drug discovery and development. Nature Reviews Drug Discovery. 2023;22:496–520.

8. Dini A, Barker H, Piki E, Sharma S, Raivola J, Murumägi A, et al. A multiplex single-cell RNA-Seq pharmacotranscriptomics pipeline for drug discovery. Nature Chemical Biology. 2024;.

9. Hsieh CY, Wen JH, Lin SM, Tseng TY, Huang JH, Huang HC, et al. scDrug: From single-cell RNA-seq to drug response prediction. Computational and Structural Biotechnology Journal. 2023;21:150–157.

10. Ding J, Adiconis X, Simmons SK, Kowalczyk MS, Hession CC, Marjanovic ND, et al. Systematic comparison of single-cell and single-nucleus RNA-sequencing methods. Nature biotechnology. 2020;38(6):737–746.

11. Wang X, Zhang R, Shen C, Kong T, Li L. Dense Contrastive Learning for Self-Supervised Visual Pre-Training; 2021.

12. Wei F, Gao Y, Wu Z, Hu H, Lin S. Aligning Pretraining for Detection via Object-Level Contrastive Learning; 2021.

13. Xie E, Ding J, Wang W, Zhan X, Xu H, Sun P, et al. DetCo: Unsupervised Contrastive Learning for Object Detection; 2021.

14. Radford A, Kim JW, Hallacy C, Ramesh A, Goh G, Agarwal S, et al. Learning Transferable Visual Models From Natural Language Supervision; 2021.

15. Jia C, Yang Y, Xia Y, Chen YT, Parekh Z, Pham H, et al. Scaling Up Visual and Vision-Language Representation Learning With Noisy Text Supervision; 2021.

16. Li Y, Liang F, Zhao L, Cui Y, Ouyang W, Shao J, et al. Supervision Exists Everywhere: A Data Efficient Contrastive Language-Image Pre-training Paradigm.; 2022.

17. Liu Y, Tian B. Protein–DNA binding sites prediction based on pre-trained protein language model and contrastive learning. Briefings in Bioinformatics. 2024;.

18. Gu Z, Luo X, Chen J, Deng M, Lai L. Hierarchical graph transformer with contrastive learning for protein function prediction. Bioinformatics. 2023;.

19. Alsaggaf I, Freitas AA, Wan C. Predicting the pro-longevity or anti-longevity effect of model organism genes with enhanced Gaussian noise augmentation-based contrastive learning on protein–protein interaction networks. NAR Genomics and Bioinformatics. 2024;.

20. Ciortan M, Defrance M. Contrastive self-supervised clustering of scRNA-seq data. BMC Bioinformatics. 2021;.

21. Yan X, Zheng R, Li M. GLOBE: a contrastive learning-based framework for integrating single-cell transcriptome datasets. Briefings in Bioinformatics. 2022;.

22. Alsaggaf I, Buchan D, Wan C. Less is more: Improving cell-type identification with augmentation-free single-cell RNA-Seq contrastive learning. Bioinformatics. 2025; p. btaf437.

23. Pedregosa F, Varoquaux G, Gramfort A, Michel V, Thirion B, Grisel O, et al. Scikit-learn: Machine learning in python. Journal of Machine Learning Research. 2011;12(85):2825–2830.

24. Chen L, Wang W, Zhai Y, Deng M. Deep soft k-means clustering with self-training for single-cell rna sequence data. NAR genomics and bioinformatics. 2020;2:lqaa039.

